# Physiological acetic acid concentrations from ethanol metabolism stimulate accumbens shell neurons via NMDAR activation in a sex-dependent manner

**DOI:** 10.1101/2023.05.05.539592

**Authors:** Andrew D. Chapp, Chinonso A. Nwakama, Paul G. Mermelstein, Mark J. Thomas

**Affiliations:** Department of Neuroscience, University of Minnesota, Minneapolis, MN 55455 USA; Medical Discovery Team on Addiction, University of Minnesota, MN 55445 USA

## Abstract

Recent studies have implicated the ethanol metabolite, acetic acid, as neuroactive, perhaps even more so than ethanol itself. In this study, we investigated sex-specific metabolism of ethanol (1, 2, and 4g/kg) to acetic acid *in vivo* to guide electrophysiology experiments in the accumbens shell (NAcSh), a key node in the mammalian reward circuit. There was a sex-dependent difference in serum acetate production, quantified via ion chromatography only at the lowest dose of ethanol (males>females). *Ex vivo* electrophysiology recordings of NAcSh neurons in brain slices demonstrated that physiological concentrations of acetic acid (2 mM and 4 mM) increased NAcSh neuronal excitability in both sexes. *N*-methyl-*D*-aspartate receptor (NMDAR) antagonists, AP5, and memantine robustly attenuated the acetic acid-induced increase in excitability. Acetic acid-induced NMDAR-dependent inward currents were greater in females compared to males. These findings suggest a novel NMDAR-dependent mechanism by which the ethanol metabolite, acetic acid, may influence neurophysiological effects in a key reward circuit in the brain.

## Introduction

Acetic acid is a short-chain fatty acid and a major metabolite of ethanol [1]. Research has recently identified acetic acid as a novel candidate mediator in the behavioral [2-5], electrophysiological [4,6,7] and epigenetic changes involved in alcohol use [8]; however, potential mechanisms of action remain scarce. Sex-specific effects in acetic acid production and mechanism of action have also yet to be investigated. With an increased prevalence of alcohol use seen in females, the sex difference in alcohol use continues to narrow [9,10]. Moreover, females demonstrate higher sensitivity to alcohol and are more likely to develop alcohol-related neuronal pathologies [9,10]. Thus, identifying potential neuroactive compounds and mechanisms are crucial to understanding alcohol use disorder (AUD) and its associated sex disparities.

Notably, ethanol is metabolized (Fig 1A) to an intermediate, acetaldehyde, primarily through the enzyme alcohol dehydrogenase. Secondary alcohol metabolizing enzymes such as catalase [1] and cytochrome P450 [1] are also capable of converting ethanol to acetaldehyde. The volatile acetaldehyde is then rapidly converted to acetic acid via aldehyde dehydrogenase. Acetic acid loses its acidic hydrogen primarily via the bicarbonate buffering system to form acetate, which can be measured with ion chromatography [11]. Alcohol research focusing on behavioral and electrophysiological effects of ethanol delivered via oral, inhaled or intraperitoneal routes often neglects this metabolic pathway and assumes that ethanol is the compound driving behavioral and neurophysiological responses. This potential oversight has left a gap in the research field, as acetic acid is a reactive and bioactive metabolite capable of disrupting pH homeostasis and influencing neuronal signaling [5,8,12-15].

**Fig 1.**
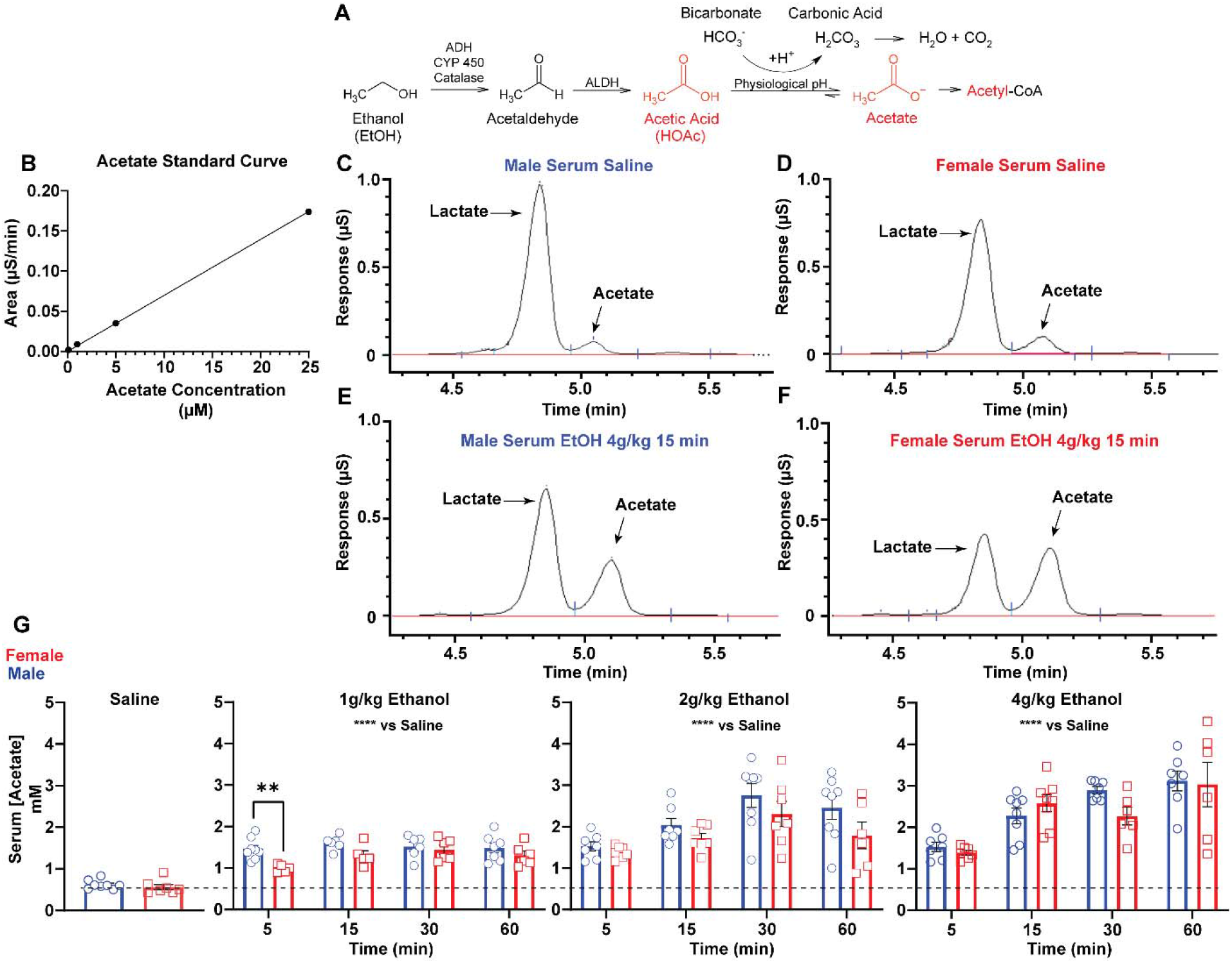
Serum acetate after ethanol exposure. **(A)** Major metabolic pathway of ethanol. **(B)** Standard curve for acetate obtained from ion chromatography (IC). **(C)** Representative IC chromatogram for male serum injected with saline depicting lactate and acetate peak separation with acetate elution time ∼5.05 min. **(D)** Representative IC chromatogram for female serum injected with saline depicting lactate and acetate peak separation with acetate elution time ∼5.05 min. **(E)** Representative IC chromatogram for male serum injected with EtOH (4g/kg) depicting lactate and acetate peak separation with acetate elution time ∼5.05 min. **(F)** Representative IC chromatogram for female serum injected with EtOH (4g/kg) depicting lactate and acetate peak separation with acetate elution time ∼5.05 min. **(G)** Summary serum acetate data of time course ethanol metabolism for male and female animals (**p=0.0046). Abbreviations: Alcohol dehydrogenase (ADH), aldehyde dehydrogenase (ALDH), cytochrome P450 (CYP 450). Dashed line depicts i.p. saline acetate levels.

To address this, we first used ion chromatography [11] to quantify serum acetic acid/acetate production and potential sex differences following acute ethanol exposure (1, 2, and 4g/kg). This was also used to inform electrophysiology studies examining the physiological effects of acetic acid on NAcSh neurons. Neurons of the NAcSh were selected for study because the direct actions of ethanol in this brain region have been reported [16], and these actions have been hypothesized to influence the rewarding aspects of this compound. We also explored the potential mechanism by which acetic acid impacted NAcSh excitability, and if there were any inherent sex differences in the acetic acid-induced response.

## MATERIALS AND METHODS

### Animals

Animal procedures were performed at the University of Minnesota Twin Cities in facilities accredited by the Association for Assessment and Accreditation of Laboratory Animal Care (AAALAC) and in accordance with protocols approved by the University of Minnesota Twin Cities Institutional Animal Care and Use Committee (IACUC), as well as the principles outlined in the National Institutes of Health *Guide for the Care and Use of Laboratory Animals*. C57BL/6J mice were obtained from The Jackson Laboratory (Bar Harbor, ME, USA), group-housed, and kept on a 14:10 light:dark cycle with food and water ad libitum.

### Chemicals

All chemicals were obtained from Sigma-Aldrich (St Louis, MO, USA) except TTX (Abcam, Boston, MA, USA), *D*-2-amino-5-phosphonovalerate (D-AP5) and memantine HCL (Cayman), and KOH IC cartridge (ThermoFisher). **Serum**

### sample collection

Male and female C57BL/6J mice (>8 weeks old) were massed and administered a single intraperitoneal (i.p.) injection of 200 proof USP standard ethanol at a dose of 1, 2 or 4g/kg body weight or an equivalent saline volume. Prior to sample collection, an i.p. injection of pentobarbital (50mg/kg) was administered to anesthetize the mice. Mice were sacrificed at the time points of 5, 15, 30 and 60 minutes after ethanol administration. To obtain estrous cycle information, vaginal cytology was performed on female C57BL/6J mice prior to serum collection.

Venous blood was obtained via needle draw from the right ventricle and deposited into 1.5 mL centrifuge tubes. Blood samples were centrifuged at 10,000 RPM for 5 minutes. The serum was removed and added to a new 1.5 mL centrifuge tube. All serum samples were placed in a -20 °C freezer until ion chromatography analysis was performed.

Serum samples were diluted 700-fold by adding 10 μL liquid sample to 6.990 mL of ddH2O (>18 MΩ) in sterile 10 mL centrifuge tubes. The samples were then transferred to ion chromatography vials (5 mL, Thermofisher) and analyzed via ion chromatography (Dionex Integrion, Thermofisher) as previously described [11].

### Quantification of acetate using IC

Diluted samples were loaded into a Dionex AS-DV autosampler (Thermofisher) connected to a Dionex Integrion RFIC system (Thermofisher) by high pressure tubing. 1 mL of sample was injected from the poly vial which was passed to the IC system. The Dionex Integrion was equipped with a Dionex EGC potassium hydroxide (KOH) RFIC, eluent generator cartridge (Thermofisher) and an AS17-C 4 mm analytical and guard column set. Water (>18 MΩ) used for generating the eluent was auto degassed within the IC system. The sample was eluted with KOH using the following method:

1. -5–0 min: Equilibration at 1 mmol/L KOH.
2. 0–10 min: Isocratic at 1 mmol/L KOH.
3. 10-17 min: Isocratic, 40 mmol/L KOH.

To determine the acetate concentration of the samples, a standard curve was constructed for known concentrations of acetate and linear regression was obtained from the areas under the curve for the known concentrations of acetate. Unknown sample acetate concentrations were determined based on the linear fit and then back-calculated based on the 700-fold dilution. Blanks subjected to the same preparation as samples were used during each batch run to assess for any detectable background acetate contamination and there was none.

### Whole-cell recordings

Mice (8-14 weeks old) were anesthetized with isoflurane (3% in O_2_) and decapitated. The brain was rapidly removed and chilled in ice cold cutting solution, containing (in mM): 228 sucrose, 2.5 KCl, 7 MgSO_4_, 1.0 NaH_2_PO_4_, 26 NaHCO_3_, 0.5 CaCl_2_, 11 d-glucose, pH 7.3-7.4, continuously gassed with 95:5 O_2_:CO_2_ to maintain pH and pO_2_. A brain block including the NAcSh region was cut and affixed to a vibrating microtome (Leica VT 1000S; Leica, Nussloch, Germany). Sagittal sections of 240 μm thickness were cut, and the slices were allowed to recover for 1 hr in a modified bicarbonate/HEPES ACSF [17], continuously gassed with 95:5 O_2_:CO_2_, containing (in mM): 92 NaCl, 2.5 KCl, 1.25 NaH_2_PO_4_, 30 NaHCO_3_, 20 HEPES, 25 glucose, 5 Na-ascorbate, 2 CaCl_2_, and 2 MgSO_4_, pH 7.25-7.3. Following recovery, slices were transferred to a glass-bottomed recording chamber circulated at a rate of 2 ml min^-1^ with standard ACSF, continuously gassed with 95:5 O_2_:CO_2_, and containing (in mM): 119 NaCl, 2.5 KCl, 1.3 MgSO_4_, 1.0 NaH_2_PO_4_, 26.2 NaHCO_3_, 2.5 CaCl_2_, 11 d-glucose, and 1.0 ascorbic acid (osmolality: 295–302 mosmol L^−1^; pH 7.3-7.4). Slices were viewed through an upright microscope (Olympus) equipped with DIC optics, an infrared (IR) filter and an IR-sensitive video camera (DAGE-MTI).

Patch electrodes (Flaming/Brown P-97, Sutter Instrument, Novato, CA) were pulled from borosilicate glass capillaries with a tip resistance of 5–10 MΩ. Electrodes were filled with a solution containing (in mM) 135 K-gluconate, 10 HEPES, 0.1 EGTA, 1.0 MgCl_2_, 1.0 NaCl, 2.0 Na_2_ATP, and 0.5 Na_2_GTP (osmolality: 280–285 mosmol L^−1^; pH 7.3)[18-20]. MSNs were identified under IR-DIC based on their morphology and hyperpolarizing membrane potential (−70 to -80 mV). MSNs were voltage clamped at - 80 mV using a Multiclamp 700B amplifier (Molecular Devices), and the currents were filtered at 2 kHz and digitized at 10 kHz. Holding potentials were not corrected for the liquid junction potential. Once a GΩ seal was obtained, slight suction was applied to break into whole-cell configuration and the cell was allowed to stabilize. Stability was determined by monitoring capacitance, membrane resistance, access resistance and resting membrane potential (V_m_) [18,19,21]. Records were not corrected for a liquid junction potential of -15 mV. Cells that met the following criteria were included in the analysis: action potential amplitude ≥50 mV from threshold to peak, resting *V*_m_ negative to −64 mV, and <20% change in series resistance during the recording.

To measure NAcSh MSN neuronal excitability, V_m_ was adjusted to -80 mV by continuous negative current injection. A series of square-wave current injections was delivered in steps of +20 pA, each for a duration of 800 ms. Working concentrations of NMDAR antagonists, AP5 (60 μM) or memantine (30 μM) were diluted from stock solutions, made in ddH_2_O and were bath applied as a cocktail with acetic acid (4 mM) for excitability studies.

To examine acetic acid-induced inward currents and the role of NMDAR in accumbens shell neurons, brain slices were continuously perfused with modified Mg^2+^-free ACSF containing (in mM): 121 NaCl, 2.5 KCl, 1.0 NaH_2_PO_4_, 26.2 NaHCO_3_, 2.5 CaCl_2_, 11 d-glucose, and 1.0 ascorbic acid (osmolality: 295–302 mosmol L^−1^; pH 7.3-7.4) at a flow of 2 mL min^-1^. Slices were gassed with 95:5 O_2_:CO_2_. Tetrodotoxin (TTX, voltage gated sodium channel blocker, 0.5 μM) and picrotoxin (GABA-A blocker, 100 μM) were added into the circulating extracellular Mg^2+^-free ACSF [7]. Cells were voltage clamped at V_m_ = -80 mV and allowed to stabilize by monitoring baseline current [7]. Once cells were stable and a baseline recording in the absence of acetic acid was observed (∼3.0 min), acetic acid (4 mM) was added to the circulating bath and the cells were recorded for 9 min. At this point, washout was observed with no recovery of any neurons tested to baseline holding currents. In separate groups of neurons, acetic acid (4mM) and memantine (30 μM) were added to the circulating bath and the NMDAR current was observed. To measure total inward current, the difference was taken between the end of drug application and baseline. Note that current-clamp and voltage-clamp experiments were performed in different NAcSh neurons.

### Statistical analysis

Data values were reported as mean ± SEM. Depending on the experiments, group means were compared using a paired Student’s *t*-test or a one-way or two-way ANOVA with repeated measures. Differences between means were considered significant at p < 0.05. Where differences were found, Bonferroni post hoc tests were used for multiple pair-wise comparisons. All statistical analyses were performed with a commercially available statistical package (GraphPad Prism, version 9.3).

## Results

### Measurement of acetic acid/acetate in serum following acute ethanol exposure

C57BL/6J mice were injected with either saline or ethanol (1, 2, or 4g/kg). Serum acetate concentrations were quantified 5, 15, 30, and 60 min post injection, using previously developed ion chromatography methodology [11]. Standard curves for acetate were linear and well fit (r^2^=0.9992) (Fig 1B), and acetate peaks displayed good resolution and consistent retention times (∼5.05 min) in both sexes (Fig 1C,D,E,F). Female estrous cycle was monitored via vaginal cytology; no estrous-dependent effects in acetate production were observed, and thus female data were pooled.

Five minutes following injection of 1g/kg ethanol, males exhibited greater acetate production than females (two-way ANOVA with Bonferroni posthoc test; 1.46 mM ± 0.09 vs 1.00 mM ± 0.043, p=0.004) (Fig 1G). At other time points and at higher doses of ethanol, no additional sex differences were observed. Ethanol increased serum acetate concentrations relative to saline at all time points tested; males (1, 2, and 4 g/kg) (one-way ANOVA, *F*_*(4,27)*_=27.58, p<0.0001, one-way ANOVA, *F*_*(4,33)*_=17.55, p<0.0001, one-way ANOVA, *F*_*(4,28)*_=36.71, p<0.0001), females (1, 2, and 4 g/kg) (one-way ANOVA, *F*_*(4,24)*_=17.50, p<0.0001, one-way ANOVA, *F*_*(4,27)*_=10.24, p<0.0001, one-way ANOVA, *F*_*(4,26)*_=14.13, p<0.0001). The maximum mean serum acetate concentration was produced at the 4g/kg dose 60 minutes after ethanol administration, in both males and females; (3.06 ± 0.35 mM and 3.03 ± 0.53 mM). The absolute maximum concentration of serum acetate measured in males was 3.96 mM and females were 4.55 mM (Fig 1G).

### Acetic acid increases excitability of NAcSh neurons

Since our experimental conditions involved a 5 min bath application of compounds and females, we conducted a time-course control as well as monitored estrous cycle and performed recordings in diestrus and estrus [6,22]. We found no differences in neuronal excitability (effect of time) in both males (two-way ANOVA, *F*_*(1,9)*_=0.8353, p=0.3846) (Fig S1,A,B) and females (two-way ANOVA, *F*_*(1,8)*_=2.977, p=0.1228) (Fig S1,C,D). Similarly, we observed no changes in neuronal excitability during different phases of the estrus cycle (two-way ANOVA, *F*_*(1,15)*_= 0.016, p=0.9012) (Fig S1,E,F).

To explore the pharmacologic impact of physiologically relevant concentrations of acetic acid guided by our *in vivo* acute experiments (Fig 1), we first assessed for a dose-dependent response to acetic acid (2 mM and 4 mM) in both males and females. We found that a 5-min bath application of both concentrations of acetic acid significantly increased the excitability of NAcSh neurons in both sexes: males 2 mM (two-way ANOVA, *F*_*(1,10)*_= 11.12, p=0.0076) (Fig 2A,B), females 2 mM (two-way ANOVA, *F*_*(1,9)*_=11.53, p=0.0079) (Fig 2C,D). Acetic acid (4 mM) in males also increased excitability, (two-way ANOVA, *F*_*(1,13)*_=25.67, p=0.0002) (Fig 2E,F), and females, (two-way ANOVA, *F*_*(1,13)*_=26.14, p=0.0002) (Fig 2G,H). Passive membrane properties between male and female NAcSh neurons did not appear to differ (Table 1).

**Table 1.** Passive NAcSh MSN membrane properties.

| | Capacitance (pF) | $V_m$ (mV) |
| --- | --- | --- |
| Male (n=54) | $83.7 \pm 2.4$ | $-76.3 \pm 0.6$ |
| Female (n=53) | $84.6 \pm 1.8$ | $-77.4 \pm 0.5$ |
| P value | 0.76 | 0.18 |

Abbreviations: $V_m$ = Resting membrane potential, $R_m$ = membrane resistance.

**Fig 2.**
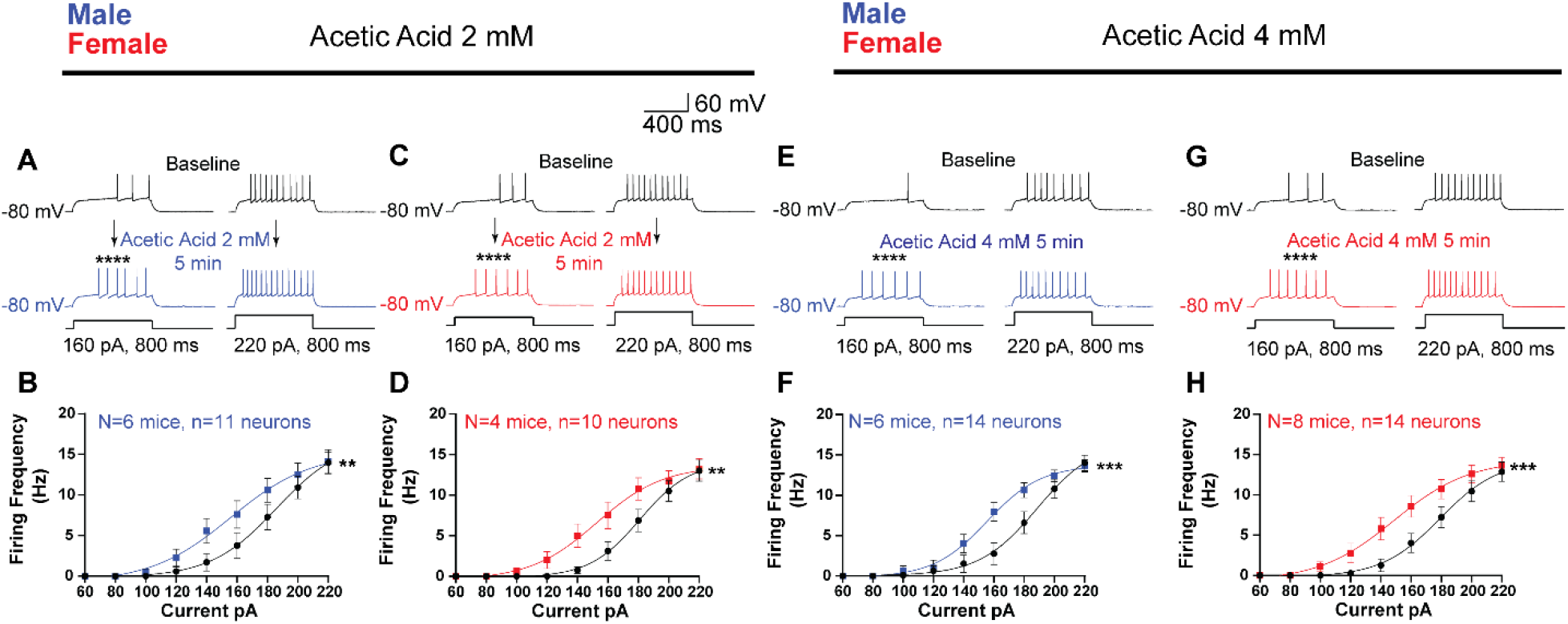
Acetic acid increases NAcSh neuronal excitability. **(A)** AP trains at 160 and 220 pA for male NAcSh neurons during acetic acid (2 mM) treatment (black, baseline; blue, 5 min after). **(B)** Summary data for current-injection response to acetic acid (2 mM) treatment for males (**p=0.0076, ****p<0.0001). **(C)** Representative traces of AP trains at 160 and 220 pA for female NAcSh neurons during acetic acid (2 mM) treatment (black, baseline; red, 5 min after). **(D)** Summary data for current-injection response to acetic acid (2 mM) treatment for females (**p=0.0079, ****p<0.0001). **(E)** AP trains at 160 and 220 pA for male NAcSh neurons during acetic acid (4 mM) treatment (black, baseline; blue, 5 min after). **(F)** Summary data for current-injection response to acetic acid (4 mM) treatment for males (***p=0.0002, ****p<0.0001). **(G)** Representative traces of AP trains at 160 and 220 pA for female NAcSh neurons during acetic acid (4 mM) treatment (black, baseline; red, 5 min after). **(H)** Summary data for current-injection response to acetic acid (4 mM) treatment for females (***p=0.0002,****p<0.0001).

### NMDAR antagonists attenuate acetic acid induced increases in NAcSh excitability

As opposed to some previous studies [23,24], here we assessed NAcSh excitability with excitatory synaptic transmission intact as we have done previously [6,19]. Thus, it remained possible that excitatory neurotransmission, perhaps via NMDARs [25], could play a role in the excitability enhancement by acetic acid. To test this, we used two NMDAR antagonists: AP5, the classical competitive antagonist and memantine, an NMDAR pore blocking antagonist. In males, the co-application of AP5 (60 μM) with acetic acid (4 mM) attenuated the acetic acid-induced increase in excitability across the stimulus response curve (two-way ANOVA, *F*_*(1,9)*_*=*2.825, p=0.1271) (Fig S2 A,B). In females, this treatment drastically blunted the effects of acetic acid across the stimulus response curve but was still significantly different from baseline (two-way ANOVA, *F*_*(1,10)*_=6.98, p=0.0246) (Fig S2 C,D). Co-application of memantine (30 μM) with acetic acid (4 mM) was able to abolish the acetic acid-induced increase in excitability across the stimulus response curve in both males (two-way ANOVA, *F*_*(1,8)*_*=*2.441, p=0.1568) (Fig 3A,B) and females (two-way ANOVA, *F*_*(1,8)*_*=*3.048, p=0.119) (Fig 3C,D).

**Fig 3.**
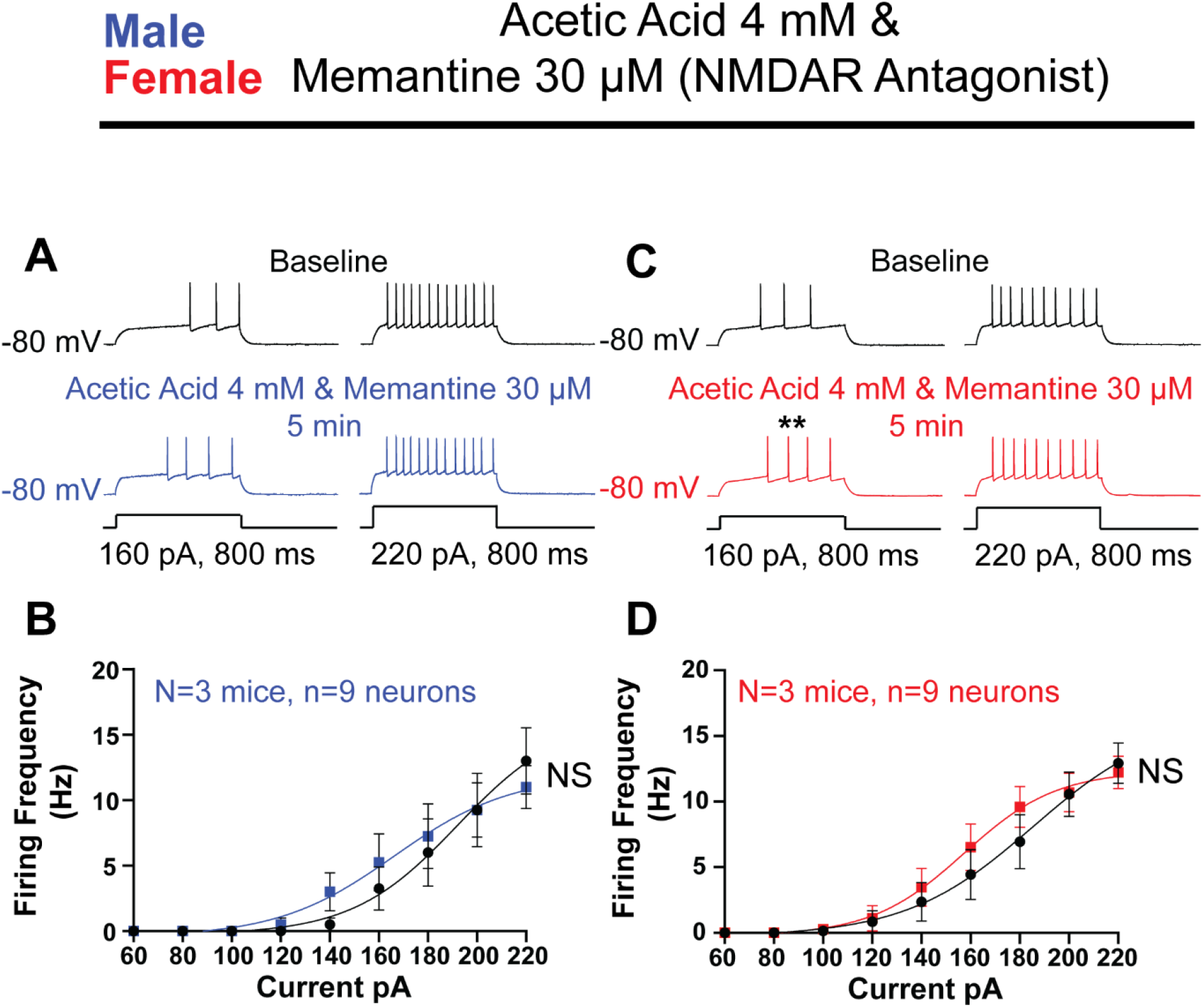
Impact of NMDAR antagonist on acetic acid induced increase in NAcSh excitability. **(A)** AP trains at 160 and 220 pA for male NAcSh neurons during acetic acid (4 mM) and memantine (30 μM) treatment (black, baseline; blue, 5 min after). **(B)** Summary data for current-injection response to acetic acid (4 mM) and memantine (30 μM) treatment for males. **(C)** Representative traces of AP trains at 160 and 220 pA for female NAcSh neurons during acetic acid (4 mM) and memantine (30 μM) treatment (black, baseline; red, 5 min after). **(D)** Summary data for current-injection response to acetic acid (4 mM) and memantine (30 μM) treatment for females (**p=0.0023).

### Acetic acid induces greater NMDAR-mediated inward currents in females than males

Given the apparent influence of NMDAR activation on the acetic acid-induced increase in excitability, we pharmacologically isolated NMDAR currents in NAcSh neurons to test for direct effects. Recording in voltage clamp mode (−80mV) in Mg^2+-^free ACSF, we found that acetic acid (4 mM) induced significant inward currents in both male (−17.38 ± 2.77 pA) and female (−54.69 ± 16 pA) NAcSh neurons compared to their baseline holding current (Fig 4). These acetic acid-induced currents were greater in females compared to males (Fig 4E, p=0.0276, two-tailed unpaired t-test). To confirm the source of these currents, we co-applied memantine (30 μM) with acetic acid (4 mM), which significantly blunted the acetic acid-elicited current in males (−17.38 ± 2.77 pA vs -7.78 ± 3.23 pA, p=0.0385, two-tailed unpaired t-test) and females (−54.69 ± 16 pA vs - 6.23 ± 5.95 pA, p=0.0131, two-tailed unpaired t-test) (Fig 4C,D), confirming its source as NMDARs. In addition to the increase in peak response (Fig. 4E), the onset kinetics of the acetic acid-induced NMDAR-mediated inward current, measured as the slope of the inward current from beginning to end of the bath application, was greater in females compared to males (−1.04×10^−4^ ± 3.44×10^−5^ pA/ms vs -3.11×10^−5^ ± 4.80×10^−5^ pA/ms, p=0.040, two-tailed unpaired t-test) (Fig 4F).

**Fig 4.**
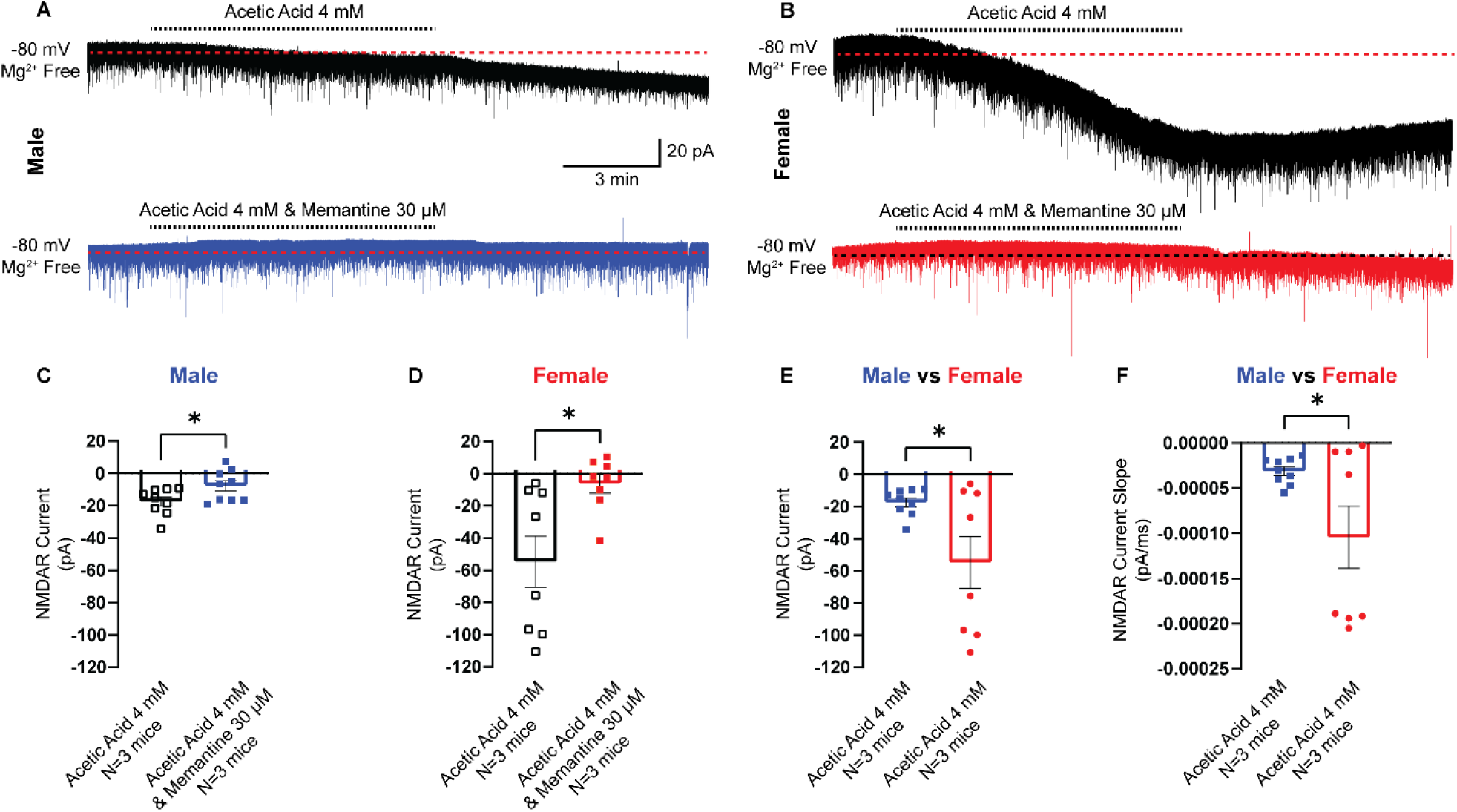
Acetic acid induces NMDAR-mediated inward currents and produces a more robust response in females. **(A)** Representative NMDAR-mediated inward current traces in male NAcSh neurons (black, acetic acid; blue, acetic acid and memantine). **(B)** Representative NMDAR-mediated inward current traces in female NAcSh neurons (black, acetic acid; red, acetic acid and memantine). **(C)** Summary data for the effects of acetic acid with and without memantine on NMDAR-mediated inward currents in male NAcSh neurons (*p= 0.0385). **(D)** Summary data for the effects of acetic acid with and without memantine on NMDAR-mediated inward currents in female NAcSh neurons (*p=0.0131). **(E)** Male vs female acetic acid-induced NMDAR currents (*p=0.0276). **(F)** Male vs female onset kinetics of acetic acid-induced NMDAR currents (*p=0.04).

## Discussion

Our study reveals several findings of relevance to alcohol research. First, we quantified serum acetic acid/acetate levels across sexes after acute ethanol exposure. At lower ethanol doses, males have a modest increased production of serum acetate compared to females. Interestingly, this sex difference is lost at higher ethanol doses. While these data provide a snapshot of possible brain concentrations of post-ethanol acetic acid levels, brain specific metabolism and/or sequestration kinetics of acetic acid/acetate via monocarboxylate transporters [26] still remains to be explored.

We also found that physiologically relevant concentrations of acetic acid (2 and 4 mM) produced robust boosts in excitability of medium spiny neurons (MSNs) in the NAcSh. This boost occurs within 5 minutes as previously reported [6], indicating the potential of acetic acid to rapidly modulate reward circuitry. Similar effects of acetate boosts to excitability have been reported in the central nucleus of the amygdala [7], suggesting that this response is likely not brain region-specific.

In addition, we determined that the effects of acetic acid on excitability were sensitive to NMDAR blockade. Interestingly, this would appear to oppose the action of ethanol at NMDARs [27]. Ethanol is widely regarded as an NMDAR antagonist [27-29], whereas our current data and previous studies provide evidence for acetic acid as an NMDAR enhancer [2,4,7]. One previous study demonstrated that high ethanol concentrations (44 mM) had no effect on NAcSh neuronal excitability, but a combination of ethanol (44 mM) and acetic acid (4 mM) increased excitability [6]. This highlights the possibility that acetic acid can override the effects of ethanol, which may be especially important *in vivo* where both ethanol and acetate are in circulation simultaneously, with acetate remaining elevated long after ethanol has been cleared [30]. Future work investigating this mixed pharmacological activity of ethanol and its metabolites at NMDARs *in vivo* will be needed to understand the neurobiological mechanisms engaged by alcohol use.

While our results implicate NMDARs in acetic acid’s effect on excitability, other mechanisms cannot yet be ruled out. For example, evidence from genetically modified mice suggests that acetate is a likely causative agent in alcohol behavioral intoxication [3,5], and this may be further supported by findings suggesting that intoxication is at least also partially mediated through acid sensing ion channels (ASICs) [31]. While this ethanol/ASICs mechanism was attributed to ethanol-hydrogen bonding interactions with the ion channel [31], we speculate the acidic hydrogen of acetic acid generated from ethanol metabolism may also be involved (Fig 1). Thus, in addition to an acetic acid/NMDAR mechanism, there seems likely to be engagement of ASICs via acetic acid. Our data demonstrate that both AP5 and memantine reduce the excitatory effects of acetic acid, although memantine was more effective than AP5 in females (Fig S2). Interestingly, memantine is also known to antagonize ASICs, specifically ASIC1A [32], the same ion channel reported to be responsible for behavioral alcohol intoxication [31]. Thus, acetic acid may activate both NMDARs and ASICs, with memantine providing more pharmacological coverage of impacted ion channels [4,7,12,32,33]. As such, we cannot rule out an acetic acid/ASICs effect within the NAcSh. However, given the significant blunting with AP5 suggests that any acetic acid-ASICs mediated excitation may be a minor effect, especially in males.

An intriguing aspect of our data is the striking sex difference in NMDAR engagement by acetic acid (Fig. 4). On one hand, this appears to follow a trend across the literature of greater NMDAR effects in females than in males [34-37]. On the other hand, given our findings of a lack of sex difference in the boost of excitability by acetic acid (Fig. 2) and a blockade of this boost by NMDAR antagonists (Fig. 3), this robust sex difference in engagement of NMDARs seems surprising. One important note in this regard, though, is that our excitability experiments were conducted with both excitatory and inhibitory synaptic transmission intact. While NAcSh synaptic neurophysiology has been well mapped in male C57BL/6J mice [38-41], the same cannot be said for females. It is possible there is stronger GABAergic tone in the female NAcSh which is only unmasked by the addition of picrotoxin—present during our NMDAR experiments, but not while assessing depolarization-induced firing. Acetate has been found to activate GABAergic signaling [3] and would be anticipated to potentially reduce some of the excitatory effects of acetic acid on NMDAR in females if they had stronger GABAergic inputs. Given the complex nature of acetic acid on excitatory [2,4,7,12,14] and inhibitory ion channels [3,14,42], further investigation of synaptic mechanisms altered by acetic acid seems warranted. Our study highlights a need for continued exploration of the effects of the ethanol metabolite, acetic acid, on behavior and neurophysiology to address the neurobiological underpinnings of Alcohol Use Disorder.

## Author Contributions

A.D.C., C.A.N., C.H.P and P.P.J performed experiments; A.D.C., C.A.N, analyzed data; A.D.C., C.A.N., P.G.M., M.J.T., prepared figures; A.D.C., C.A.N., P.G.M., and M.J.T. drafted manuscript; A.D.C., C.A.N., P.G.M. and M.J.T. interpreted results of experiment; A.D.C., C.A.N., P.G.M. and M.J.T. edited and revised manuscript.

## Acknowledgements

We would like to thank Chau-Mi H. Phan and Pramit J. Jagtap for their assistance. We would also like to thank, Dr. Andréa R. Collins, Dr. Timothy W. Chapp, Dr. Scott M. Chapp, Casey L. Mallo and Dr. Qing-Hui Chen for their proofreading and suggestions.

## Conflicts of interest

The authors declare no conflicts of interest.

### Supplemental Data

**Fig S1.**
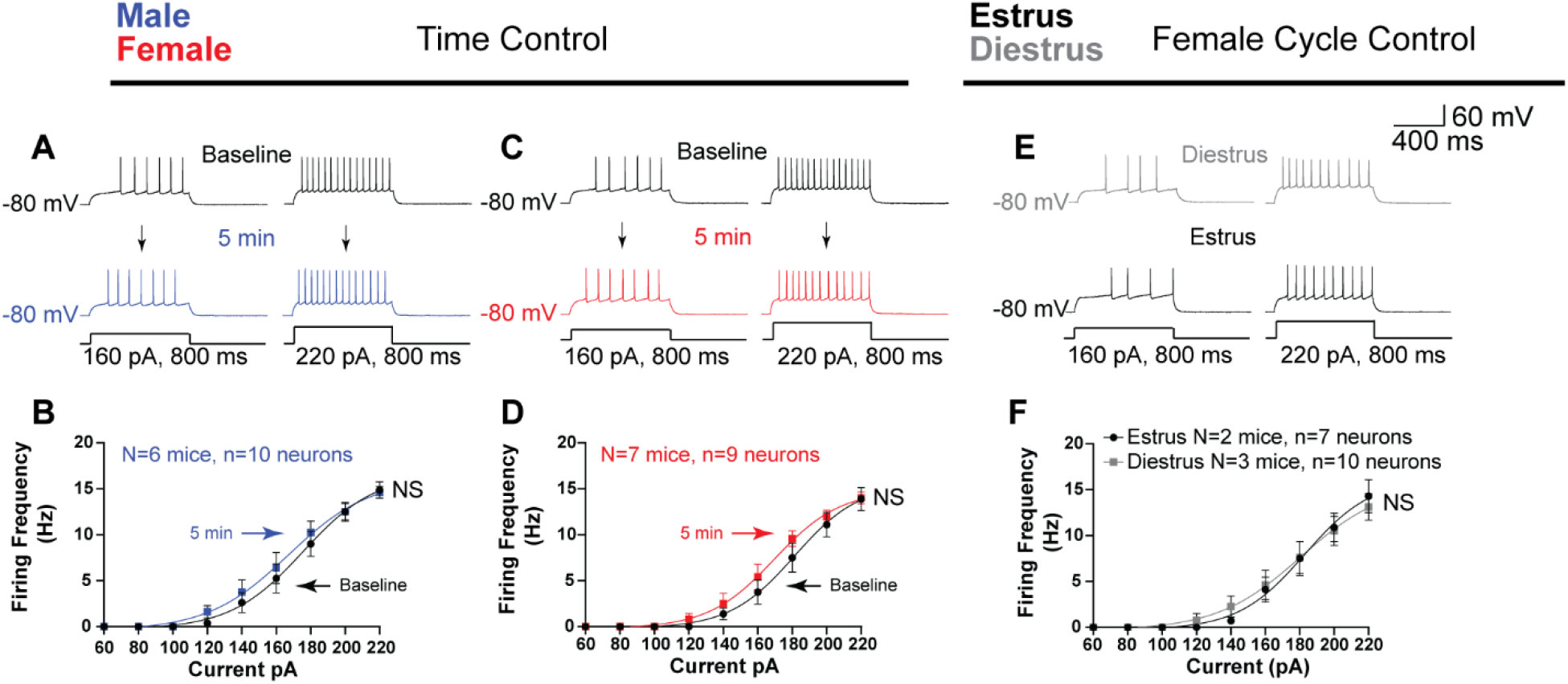
Effect of time and estrous cycle on NAcSh neuronal excitability. **(A)** Representative raw traces from male accumbens shell neurons at baseline (black, top) and after 5 min holding (bottom, blue). **(B)** Summary male data of stimulus response for time course control. **(C)** Representative raw traces from female accumbens shell neurons at baseline (black, top) and after 5 min holding (bottom, red). **(B)** Summary female data of stimulus response for time course control. **(E)** Representative raw traces from female accumbens shell neurons at 160 pA (left) and 220 pA (right) during diestrus (gray) and estrus (black). **(F)** Summary data for the current injection response of female accumbens shell neurons in diestrus (gray) and estrus (black). NS=not significant.

**Fig S2.**
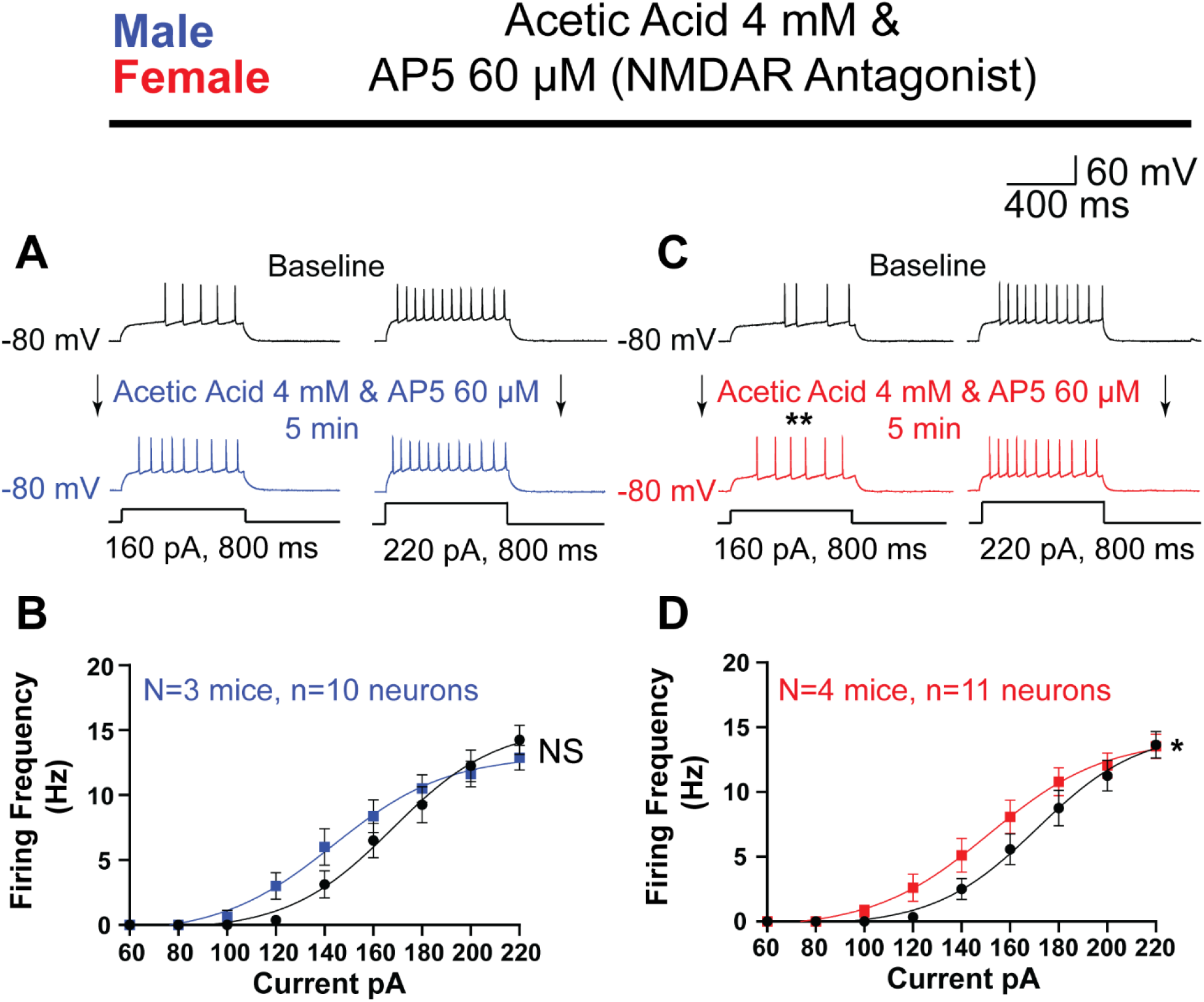
Impact of AP5 on acetic acid induced increase in NAcSh excitability. **(A)** AP trains at 160 and 220 pA for male NAcSh neurons during acetic acid (4 mM) and AP5 (60 μM) treatment (black, baseline; blue, 5 min after). **(B)** Summary data for current-injection response to acetic acid (4 mM) and AP5 (60 μM) treatment for males. **(C)** Representative traces of AP trains at 160 and 220 pA for female NAcSh neurons during acetic acid (4 mM) and AP5 (60 μM) treatment (black, baseline; red, 5 min after). **(D)** Summary data for current-injection response to acetic acid (4 mM) and AP5 (60 μM) treatment for females (*p=0.0246,**p=0.0016).

